# Syncytial nerve net in a ctenophore sheds new light on the early evolution of nervous systems

**DOI:** 10.1101/2022.08.14.503905

**Authors:** Pawel Burkhardt, Astrid Medhus, Leonid Digel, Benjamin Naumann, Joan J Soto-Àngel, Eva-Lena Nordmann, Maria Y Sachkova, Maike Kittelmann

## Abstract

A fundamental breakthrough in neurobiology has been the formulation of the neuron doctrine by Santiago Ramón y Cajal, which states that the nervous system is composed of discrete individual cells. Electron microscopy later confirmed the doctrine and allowed the identification of synaptic connections. Here we use volume electron microscopy and 3D reconstructions to characterize the nerve net of a cydippid-phase ctenophore, belonging to one of the earliest-branching animal lineages. We found that neurons of its subepithelial nerve net do not follow Cajal’s neuron doctrine but instead show a continuous plasma membrane forming a syncytium. This is more similar to the reticulate theory of the nervous system put forward by Camillo Golgi. Additionally, we were able to identify new sensory cell types and describe simple neuro-sensory circuits for cydippid-phase ctenophores. Together with the ctenophore-specific synaptic architecture and the presence of an extensive repertoire of lineage-specific neuropeptides our morphological data provide substantial evidence for the independent evolution of the nervous system of ctenophores and the remaining animals.

## Introduction

For more than one century, the structure and evolutionary origin of the animal nervous system has been at the centre of much debate among biologists. Fundamental progress in our structural understanding was put forward by Santiago Ramón y Cajal, postulating that the nervous system is composed of discrete cells, so-called neurons, rather than forming a syncytial continuum, as proposed by Camillo Golgi^1^. The discovery of synaptic connections between individual neurons by electron microscopy later confirmed Cajal’s “neuron doctrine”. But did such neurons and their organization into a nervous system evolve only once? There is accumulating evidence that ctenophores, gelatinous marine invertebrates moving through the water column by ciliary comb rows, are the sister-group to all other animals (Figure 1A)^2–5^. Ctenophores exhibit a complex life cycle including a predatory cydippid stage that hatches from the egg and is already able to reproduce after a few days (Figure 1B)^6^. Ancestral state reconstruction suggests the cydippid body plan is a plesiomorphic character of ctenophores^7^

**Figure 1.**
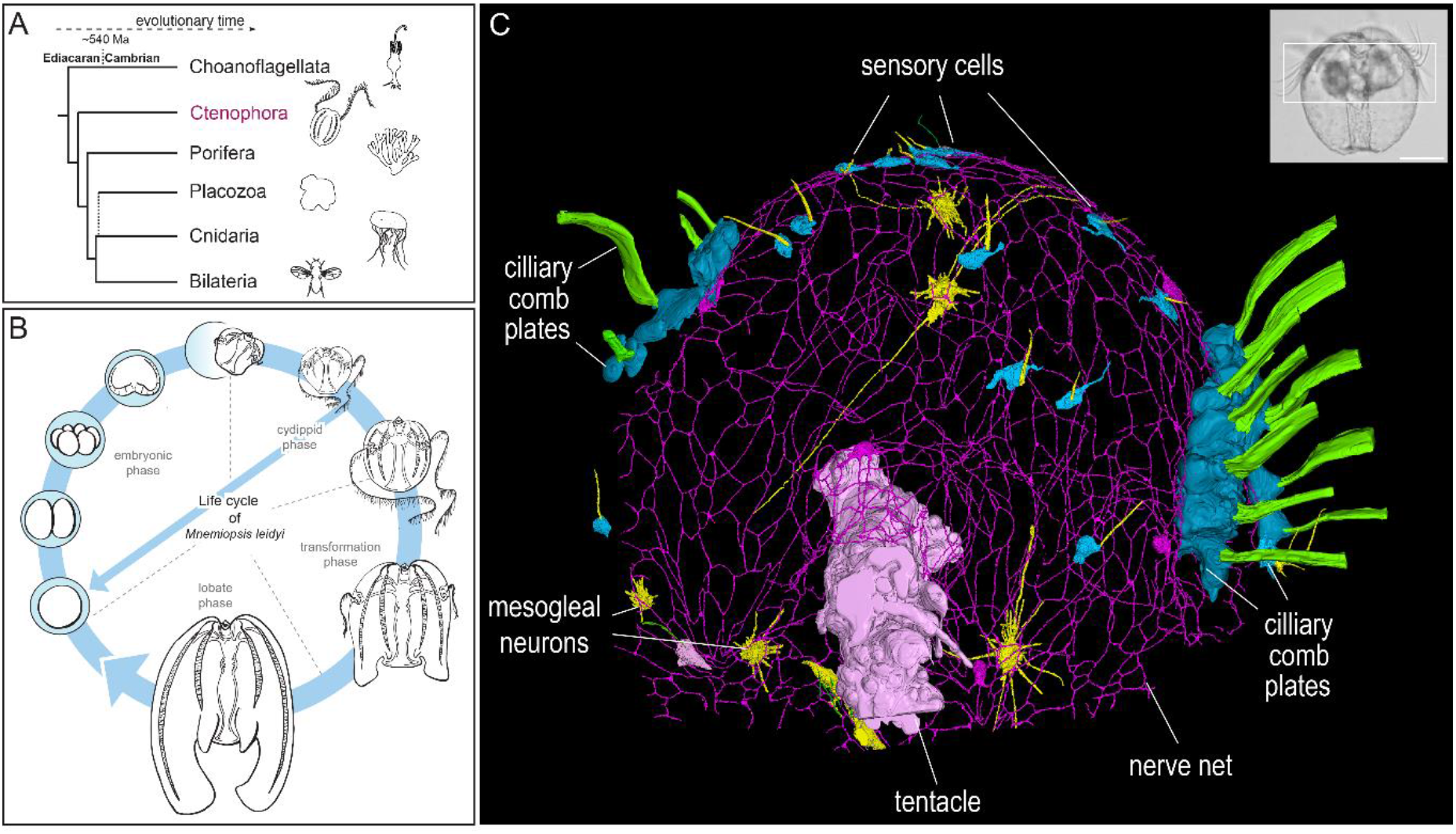
Ctenophores and their nervous system. (A) Phylogenetic placement of ctenophores within the animal tree^2,4,5^. (B) The ctenophore *Mnemiopsis leidyi* exhibits complex life cycle stages including a predatory cydippid phase that hatches from the egg and can reproduce after a few days. (C) 3D reconstruction of the nerve net, comb rows, sensory cells, mesogleal neurons and a tentacle from SBFSEM data of a 1-day old cydippid. Inset: Phase contrast image of a 1-day old cydippid. White box: area reconstructed in C. Scale bar: 100 μm.

The early split of ctenophores from other groups indicates that a nervous system, and maybe even neurons, evolved at least twice – once within the ctenophores and once within the lineage of the remaining animals^8^. Initiated by genomic analyses^2,3^, molecular and physiological features of the ctenophore nervous system were subsequently interpreted to support this scenario^4,5^. In contrast to sponges and placozoans, ctenophores exhibit an elaborate nervous system consisting of a subepithelial nerve net (SNN), mesogleal neurons, an aboral (sensory) organ, tentacle nerves and diverse sensory cells in all parts of their body (Figure 1C and Suppl. Video 1)^9–14^. A further unique feature is the structure of the ctenophore synapse: a mitochondrion, ER sheet and synaptic vesicles form a tripartite complex termed the “presynaptic triad”^15,16^. These peculiarities show that deciphering the development, structure and function of the ctenophore nervous system is a key element to understand the origin and evolution of animal nervous systems. We have recently shown that a large repertoire of lineage-specific neuropeptides has evolved in the ctenophore *Mnemiopsis leidyi*^14^. Furthermore, we identified unique SNN neurons extending multiple neurites that interconnect through anastomoses and thus form an extensive continuous network^14^. This characteristic sets them apart from other animal neurons, further supporting a possible independent evolution and hinting to an alternative neuronal organization different from Cajal’s “neuron doctrine”. To test this, it is necessary to understand how SNN neurons connect to each other, to sensory neurons and to cells within the mesoglea. Here we used high pressure freezing fixation techniques in combination with Serial Block Face Scanning Electron Microscopy (SBFSEM) to establish the first ultrastructural 3D network of SNN neurons and other cell types in a ctenophore.

## Results

### The cydippid SNN is organized in a syncytium

The relatively recent development of SBFSEM allowed the collection of large volume image data of whole-mount cydippid-phase *M. leidyi* at electron microscopy resolution. Detailed analysis of an early cydippid revealed that the neurites of all five SNN cells in the dataset were connected through an anastomosed continuous network (Figure 2A). Neither electrical nor chemical synapses were detected between the cells of the SNN. Neurites within the SNN exhibit no obvious polarity (axon vs. dendrite) regarding their morphology, showing similar diameters, dense core vesicles distribution throughout their length and the lack of the typical presynaptic triads (Figure 2B and C).

**Figure 2.**
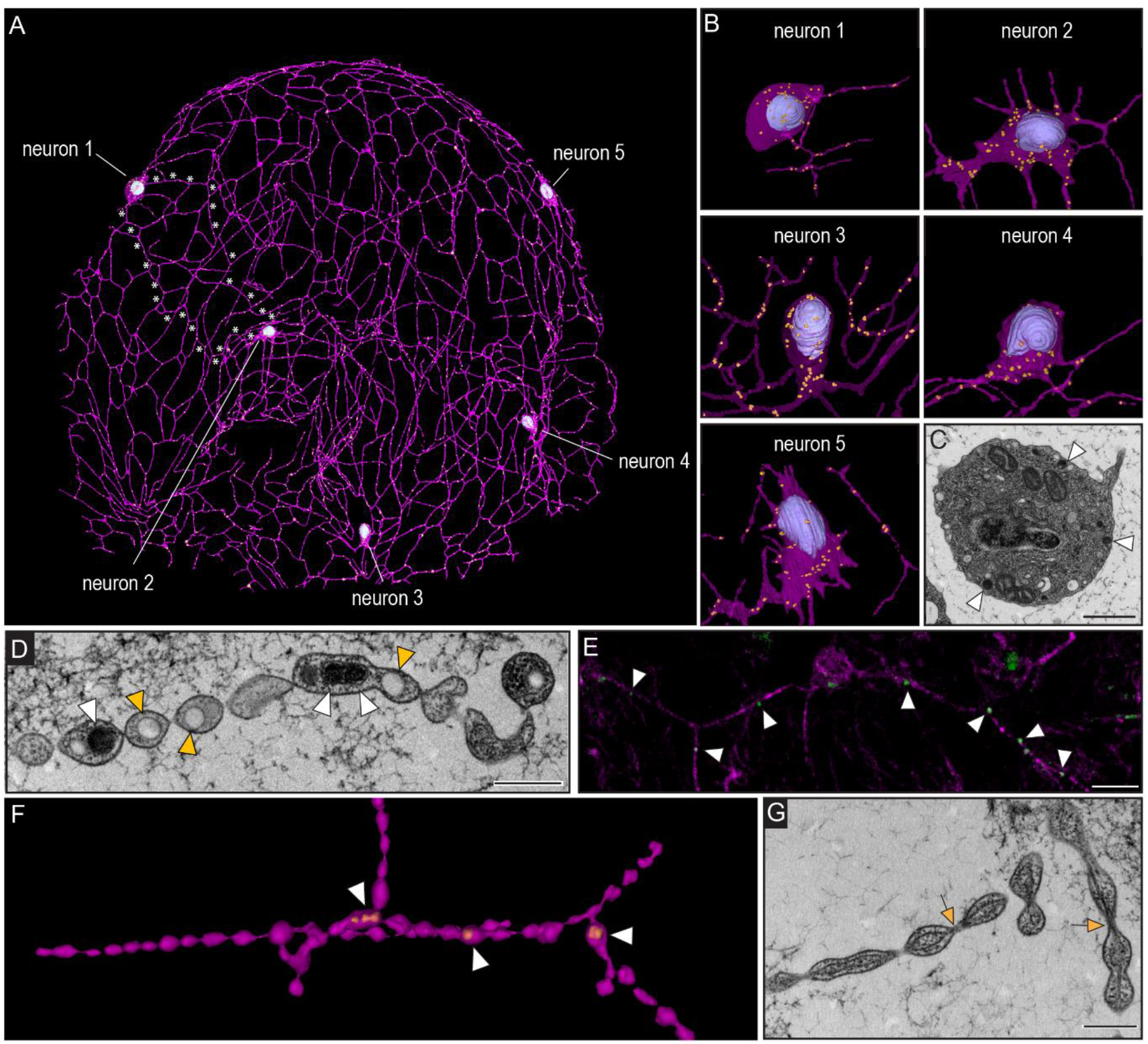
Connectivity and ultrastructure of the ctenophore SNN. (A) 3D reconstruction of five SNN neurons. White asterisks indicate examples of continues membrane between cell bodies of neuron 1 and 2. (B) 3D reconstruction of the SNN neuron cell bodies showing the nucleus (blue) and dense core vesicles (orange). (C) TEM cross section of an SNN neuron cell body showing ultrastructural details including large dense core vesicles (white arrowhead). (D) TEM cross section of a SNN neurite with dense core and clear core vesicles localized in “blebbed” areas (white and orange arrowheads). (E) Antibody staining against neuropeptide ML199816a (green) in SNN neurites (magenta) stained with anti-tubulin. (F) TEM 3D reconstruction of SNN neurite (violet) and dense core vesicles (orange) highlighting the blebbed morphology. (G) TEM cross section of SNN neurites showing continuous microtubules (orange arrows) passing through narrow segments. Scale bars C: 1 μm; D, G: 500 nm.

Strikingly, SNN neurites often showed a blebbed or “pearls-on-a-string” morphology (Figure 2D-G and Suppl. Figure 1). The narrow segments are often just wide enough for microtubules to pass (Figure 2G, Suppl. Figure 1), and bulged segments often contain larger clear or electron dense vesicles and occasionally ER (Figure 2D and Suppl. Figure 1). A newly developed antibody against the neuropeptide ML199816a^14^ confirms the presence of neuropeptides within some of the vesicles of SNN neurons (Figure 2E). While SNN neurons seem to lack synapses between each other, we identified chemical synapses from the SNN to polster cells (Suppl. Figure 2), suggesting directional signal transmission from the SNN to effector cells.

### Mesogleal neurons form direct contacts with the syncytial SNN

We identified and reconstructed six mesogleal neurons that exhibit a star-like morphology with extensive plasma membrane protrusions of variable lengths (Figure 3A). Their somata are filled with a variety of vesicles and larger vacuoles (Figure 3B). The protrusions of these cells do not show the “pearls-on-a-strings” morphology present in neurites of the SNN. Some of the protrusions form plasma membrane contacts to neurites of the SNN (Figure 3A, D, E). However, we did not find ultrastructural evidence for electrical or chemical synapses (Figure 3E). In contrast to SNN neurons, we did not observe any electron dense vesicles in mesogleal neurons (Figure 3B) but instead small electron-lucent vesicles of a similar size as synaptic vesicles (Fig. 3C). This clearly distinguishes them from SNN neurons (Figure 2) and suggests a different type of information transmission.

**Figure 3.**
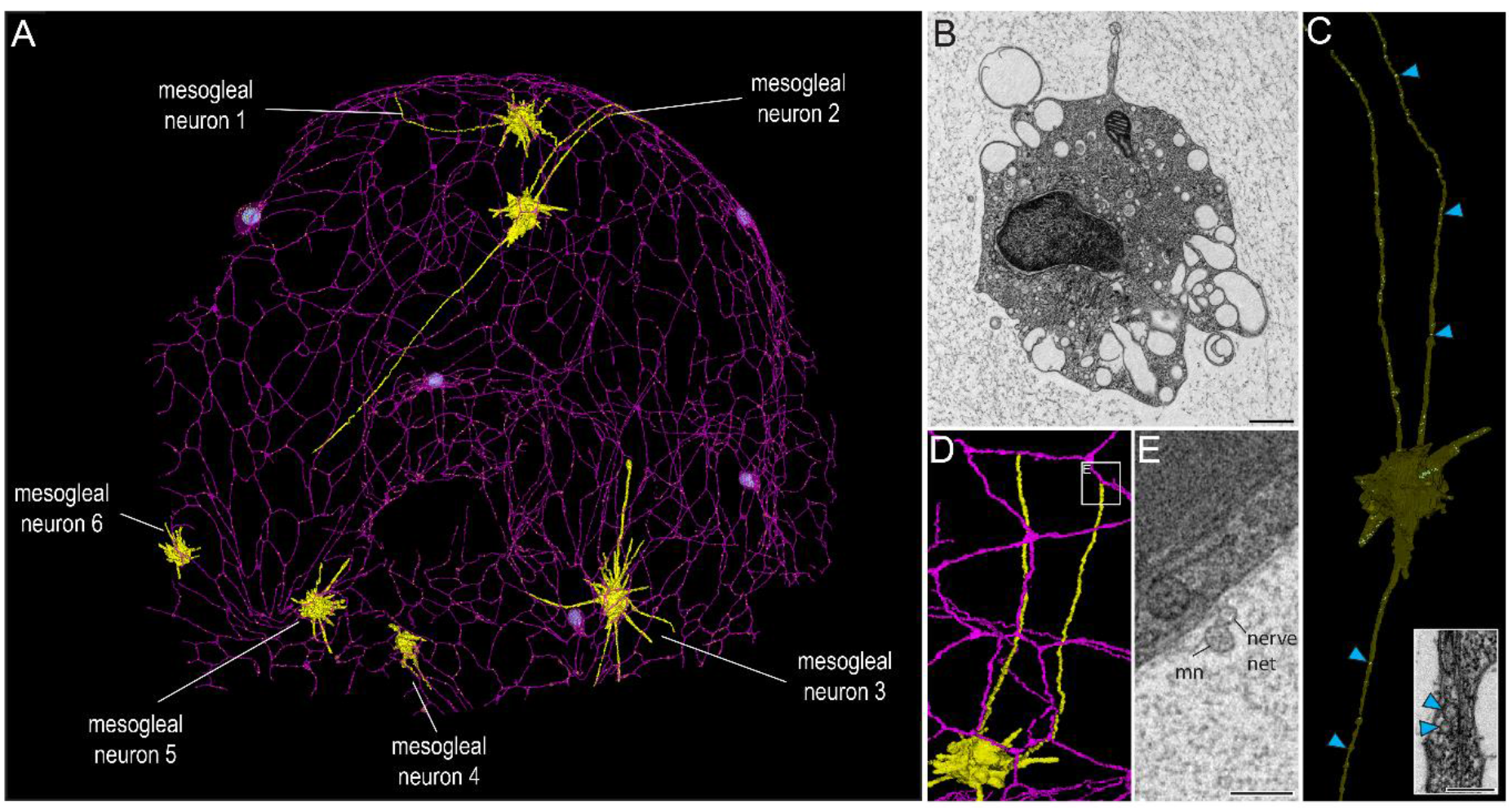
Close association of mesogleal neurons and the SNN. (A) 3D reconstruction of SNN (violet) and mesogleal neurons (yellow) from SBFSEM data. (B) TEM cross section of a mesogleal neuron cell body. Different types of clear vesicles and vacuoles but no dense core vesicles are present. (C) 3D reconstructed mesogleal neuron with three long neurites that contain small clear vesicles (blue arrowheads). TEM cross section of mesogleal neurites with small clear vesicles shown in inset. (D) 3D reconstruction of mesogleal neuron with contact site (white box) to SNN. (E) Corresponding SBFSEM image of contact site between mesogleal neuron and SNN neuron. mn: mesogleal neuron. No chemical or electric synapse structures could be observed. Scale bars B: 1 μm; C (inset): 200 nm; E: 500 nm.

### Sensory cells form simple circuits involving the syncytial SNN

We identified and reconstructed a total of 22 putative sensory cells from the present and an earlier data set published by Sachkova et al. (2021)^14^ which fit into five morphological groupings (Figure 4, Suppl. Figure 3 and Suppl. Table 1). Some of them resemble known ctenophore sensory cell types (type 1, 4 and 5)^16,17^ while others exhibit a morphology not described previously (type 2 and 3) (Figure 4, Suppl. Figure 3, and Suppl. Table 1). We detected chemical synapses in several but not all putative sensory cells contacting neuronal or other effector cells (Figure 4, Suppl. Figure 3). Type 1 sensory cells exhibit a single long cilium and onion root basal body (Figure 4, Suppl. Figure 3). Type 2 sensory cells exhibit a very short single cilium without an onion root basal body. Long neurites extending from their somata form chemical synapses to polster cells (Figure 4B, Suppl. Figure 3A and C).

**Figure 4.**
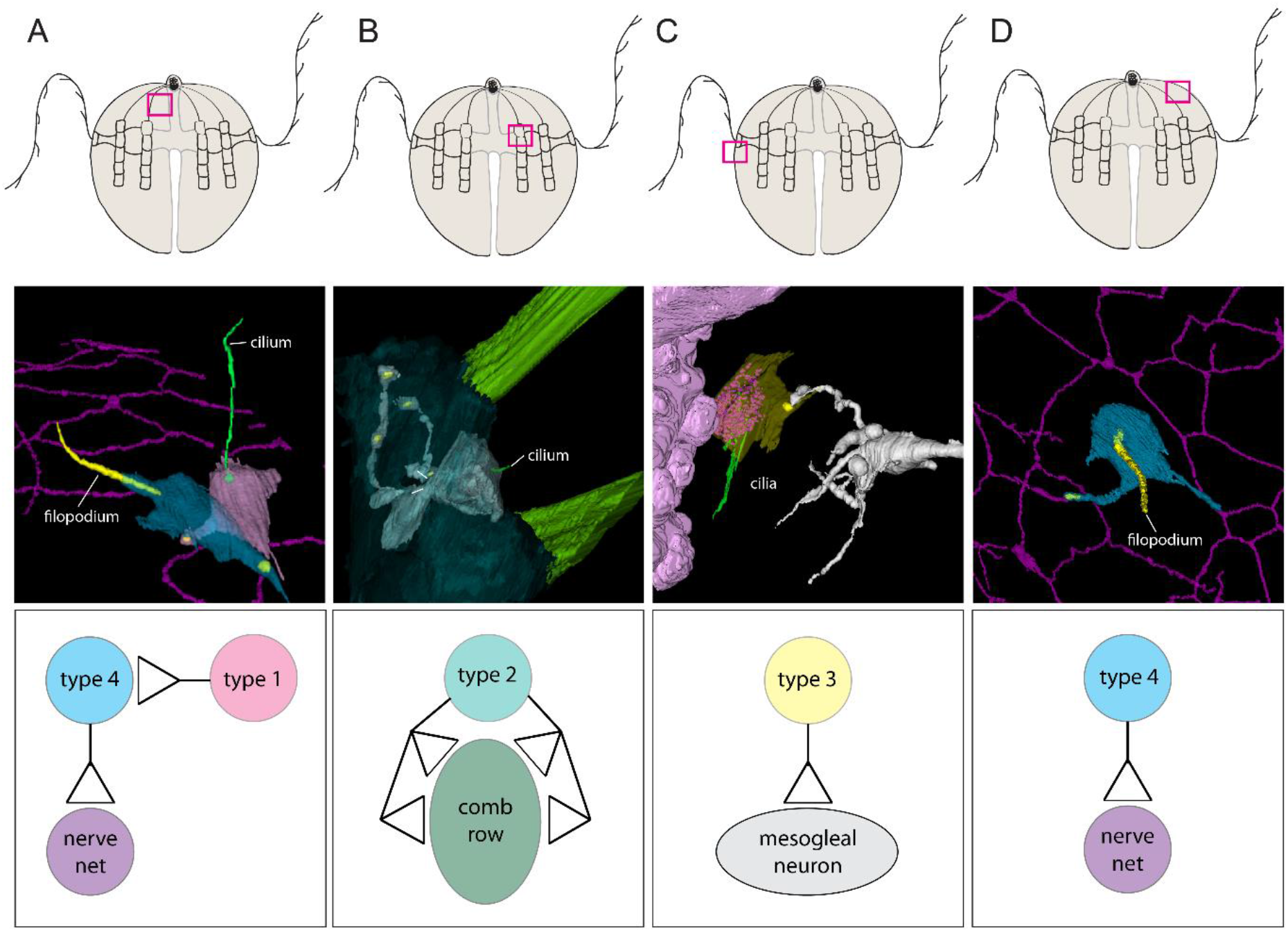
3D reconstruction of sensory cells allows for the identification of simple circuits. Top panel: Localization of each circuit (pink square). Middle panel: 3D reconstructions of sensory and effector cells. Mitochondria are shown in yellow as representative of synaptic tripartite complexes in all circuits. Bottom panel: Proposed wiring diagram. A) Circuit between type 1 and type 4 sensory cell and SNN. (B) Multiple synaptic connections between type 2 sensory cell with short cilium and comb cells. (C) Synaptic connection between type 3 sensory cell near tentacle and a mesogleal neuron. (D) Type 4 sensory cell with single filopodium synapses onto nerve net.

Type 3 sensory cells exhibit multiple cilia without onion root basal bodies. Many large electron dense vesicles are localized beneath the cilia (Figure 4C and Suppl. Figure 3A and D). We found one of these cells near the tentacle with a synaptic connection to a mesogleal neuron (Figure 4C). Type 4 sensory cells exhibit a single long filopodium. Some of them form synapses to neurites of the SNN (Figure 4A and D) and some also receive synaptic input from type 1 sensory cells (Figure 4A). Type 5 sensory cells exhibit multiple long filopodia. They can form plasma membrane contact to polster cells. We were not able to detect synaptic contacts from or to this cell type. Using SBFSEM based 3D reconstruction we are now able to propose several discrete and simple neural circuits in early cydippid-phase *M. leidyi*. These circuits include synaptic signal transmission from sensory cells to other cell types including SNN neurons, mesogleal neurons, polster cells or even other sensory cell types (Figure 4A-D).

## Discussion

In the lively debates about the organization of animal nervous system at the end of the 19^th^ century Joseph von Gerlach (1871)^18^ and Camillo Golgi (1885)^19^ put forward the “reticular theory” (also syncytial theory). Both proposed the cellular continuity of neurons. This view was challenged by the “neuron doctrine” of Ramón y Cajal (1888)^1^ proposing an organization from discrete cellular units connected via synapses. Both contestant theories were founded on Golgi’s newly invented black staining that enabled scientists to study the detailed morphology of neurons and their neurites^20^. Golgi and Cajal were honored with the Nobel Prize in Physiology or Medicine in 1906 for their effort in elucidating the architecture of the nervous system^20^. However, with the advent of electron microscopy in the 1950s and the discovery of the synaptic cleft, the reticular theory was put to rest in favor of Cajal’s neuron doctrine^21,22^. In the present study, volume electron microscopy revealed the 3D ultrastructural architecture of the SNN in an early cydippid-phase ctenophore providing evidence for its reticular – or syncytial – organization. Using high pressure freezing and freeze substitution techniques to preserve fine ultrastructural details with minimal fixation artifacts, we show that the SNN is a continuous structure that connects externally to polster and mesogleal neurons via synaptic triads and other plasma membrane contacts. Previous characterizations of ctenophore nerve nets have been predominantly based on traditional histochemical staining techniques^9,23^, and more recently on fluorescence microscopy of antibody staining against alpha-tubulin^10,12,13,24^. Both of these techniques do not allow investigating the ultrastructure of neuronal connections. Superb data from transmission electron microscopic serial sections^15,25^ may also have overlooked this special architecture due to the difficulty to produce continuous section series. Interestingly, besides reports on single self-anastomosing neurites in other animals^26–28^, the presence of a complete syncytial (“closed”) nerve net has only been reported for cnidarian, medusae-like colonial polyp *Velella*^29,30^. However, the syncytial organization of this nerve net has not yet been verified on an ultra-structural level.

While neurite fusion and pruning seem to be a common principle during the early neural development in many animals^31,32^ we do not consider the syncytial cydippid SNN do be completely remodeled by such a process. It was suggested that the early cydippid-phase is not a larval but rather autonomous life history phase of *M. leidyi* and other ctenophores^6^. Cydippid-phase *M. leidyi* are free-swimming pelagic predators, able to reproduce and exhibit complex behaviors similar to their second reproductive, lobate-phase^33–35^. Even if the syncytial organization of the SNN becomes synaptic during this cydippid-lobate-transition, *M. leidyi* offers a unique new model system to understand basic neuronal mechanisms. It can be used to investigate a new developmental process in which a functional syncytial nerve net is remodeled into a functional synaptic system or utilized to understand the development and function of a unique nervous system combining a neuronal syncytium as well as synaptic connection.

The discovery of the non-synaptic architecture of the cydippid-phase SNN raises the intriguing question about the mechanism of signal propagation. Genome and single cell transcriptome analyses revealed that *M. leidyi* SNN neurons express several voltage gated calcium (Ca_v_), 35 potassium (K_v_) and two non-specific sodium (Na_v_) channels^14,36,37^. These numbers are similar to neurons of other animals and ctenophore SNN neurons are therefore very likely to produce action potentials^38^. Additionally, the presence of numerous peptidergic vesicles in the SNN suggests that signal transmission also occurs through neuropeptide release, and Ca_v_2 channel expressed in these cells might be involved in exocytosis^14,39^. Therefore, the SNN could function as a neuroendocrine system that is able to release transmitters into the mesoglea via vesicle fusion with the plasma membrane at different neurite sites. Such a system would require only a minimum number of chemical synapses and, if acting at short distances, may reach enough effector cells. Indeed, studies on the conduction velocity in ctenophores have shown a slower speed of signal propagation compared to nerve nets and conducting epithelia of other animals^40^, indicating that signal propagation could be non-synaptic.

Additionally, our identification of simple circuits now allows for a better understanding of mechanoreception, swimming and prey capture behavior in young cydippid-phase ctenophores. Type 1 ciliated sensory cells and type 4 filopodiated sensory cells, previously described as ‘Tastborsten’ and ‘Taststifte’ by Hertwig in 1880^9^, have been postulated to be sensitive to water vibrations and touch^17,41,42^. Their abundance throughout the epidermis and direct cell-cell contact to the nerve (many through chemical synapses) highlights the importance of localized vibration and touch information to be transmitted directly to the SNN. A type 2 sensory cell, which wraps around polster cells, may be able to detect water flow and thus alter comb beat frequency whereas a type 3 sensory cell, whose multiple cilia are in close contact to the tentacle, may be triggered by food capture.

The appearance of first nerve nets is strongly interconnected with the origin of neurons and highlights the importance of studying nerve nets^43^. However, whether the neuron of animals has a single origin or possibly originated more than once during evolution is a hotly debated topic. The presence of a syncytium-like SNN highlights that ctenophores came up with a different way to build a neural network and provides tentative evidence that the ctenophore nerve net evolved independently. Our ultrastructural analysis of the ctenophore SNN not only puts ctenophores at the center of nervous system evolution, but also provides a unique opportunity to explore the boundaries of nervous system organization and function.

## Material & Methods

### Animal husbandry

For electron microscopy experiments, *M. leidyi* cydippids were 1-day old (i.e., < 24 hours after hatching or < 48 hours post fertilization). Animals were obtained from two months old, 3-5 mm cydippids as previously described^14^. Briefly, the parental generation was kept in 300 ml beakers with 5-10 individuals per beaker in sea water at 20-22 °C, 27 ppt and pH 7.9 – 8.1. Ctenophores were fed with living *Brachionus* (rotifers) once a day and five times a week with a final density of 10 prey/ml. Beakers were washed every five days by carefully transferring the cydippids into beakers with new seawater. Seawater used for both the ctenophore and the rotifer culture was first filtered through a combination of 10, 5 and 1 μm mechanical filters, activated charcoal and UV irradiation. *Brachionus* were kept in 6 L transparent buckets and fed with 8-10 ml of commercial concentrated microalgae RGcomplete™ distributed in 2-3 dose/day and five times a week. In the described conditions, cydippids become reproductive ca. one week after hatching and spawn daily and continuously for years. Hatching occurs 22-26 hours after spawning and fertilization.

### Immunohistochemistry

The mature neuropeptide deriving from the ML199816a precursor was predicted earlier^14^. The peptide was chemically synthesized with the addition of an extra Cys residue at the C-terminus (ML199816a, EEDSAFLFADC) to enable conjugation to KLH and used for immunization of rabbits followed by affinity purification by Genscript.

Animals were fixed 3-4 days post fertilization in ~ 16% Rain-X® in artificial sea water (ASW) for 1hr at room temperature (RT), followed by further fixation in ice cold 3.7% formaldehyde in ASW for 1hr on ice^44^. Cydippids were washed four times in PTW buffer (1.8 mM KH_2_PO_4_, 10 mM Na_2_HPO_4_, 0.137 M NaCl, 2.7 mM KCl, 0.1% Tween-20, pH 7.4) or until no Rain-X® droplets were observed. The cydippids were stored in PTW at 4°C up to a week. Animals stored for longer were dehydrated trough a methanol series in PTW [50%, 75%, 100% (v/v)] and kept at −20°C and subsequently rehydrated trough methanol in PTW series [60%, 30% and 0% (v/v)] before use. Animals to undergo immunostaining were washed five times for 5 minutes in PBTx (0.2% Triton X100 in PBS (137 mM NaCl, 2.68 mM KCl, 10.14 mM Na2HPO4, 1.76 mM KH_2_PO_4_, pH 7.4)), before blocking with 1% Bovine serum albumin in PBTx for 1hr at RT. ML199816a antibody was combined with mouse E7 beta tubulin antibody (DSHB). The primary antibodies were diluted 1:100 in blocking solution, spun down at 16000 rcf for 10 minutes, and the supernatant was used for overnight incubation at 4°C. Samples were subsequently washed six times in PBTx for 15 minutes at RT, before secondary antibodies, goat anti-rabbit 647 (ab150083) and goat anti-mouse 488 (ab15017), were diluted 1:250 in blocking solution, spun down at 16000 rcf for 10 min and the supernatant was used to stain the animals overnight 4°C. The samples were washed three times in PBTx for 15 min followed by five 5-minute washes in PBS and mounted in Vectashield antifade mounting medium containing DAPI (Vector Laboratories). Samples were imaged on an Olympus FV3000 Confocal Laser Scanning Microscope and processed in Imaris.

### TEM and SBFSEM sample preparation and imaging

1-day old *M. leidyi* cydippids were fixed and imaged using High Pressure Freezing and Freeze substitution as previously described^14^. Briefly, *M. leidyi* cydippids were frozen in 20% BSA in seawater (Baltech HPM010), freeze substituted with a mix of 1% UA and 1% Osmium in acetone over 72 hours from −90 to 4 °C. Additional en block staining with 1% tannic acid in acetone for 2 hrs and subsequently with 1% osmium in acetone for 1 hr was performed at room temperature. Ctenophores were then infiltrated and embedded in 812 Epoxy resin and cured at 60 °C for 30 hrs.

SBFSEM images were collected with a Merlin Compact SEM (Zeiss, Cambridge, UK) with the Gatan 3View system and Gatan OnPoint BSD with pixel size 5 nm, dwell time 1us, 20 nm Aperture, 1.8 kV acceleration voltage in high vacuum with Zeiss FocalCC set to 100%. Section thickness was 100 nm.

Series of 50 nm thin sections for TEM were collected with PowerTome ultramicrotome (RMC) and imaged with a Jeol JEM-1400Flash with a Gatan OneView 16 Megapixel camera at 120kV.

### 3D reconstruction

The dataset was binned in X and Y to a resolution of 30×30×100 nm voxel size. 3D reconstruction was performed as previously described^14^. Briefly, to reconstruct a *M. leidyi* whole mount cydippid SBFSEM sections were imported as z stacks into the Fiji^45^ plugin TrakEM2^46^ and automatically aligned using default parameters. Alignments were manually curated and adjusted if deemed unsatisfactory. Whole cells (SNN, comb cells, mesogleal neurons, sensory cells), tentacle and organelles were manually segmented, and 3D reconstructed by automatically merging traced features. Meshes were then smoothed in TrakEM2. For the generation of the 3D animation the .obj file from TrakEM2 was exported and imported into Blender 3.0.1^47^. The membranes of the SNN were rendered transparent to reveal the underlying dense core vesicles. No other transformations that could affect the cells’ morphology have been performed. The entire EM reconstruction was animated using the basic suite of 3D animation functions of the software at 24 frames per second.

## Author contributions

PB and MK designed the study; PB, AM, JJSA, ELN, MS, and MK performed experiments; LD generated the 3D video animation. PB, AM, MS and MK analyzed data; PB, BN and MK wrote the paper and all authors reviewed, commented on, and edited the manuscript.

## Acknowledgments

The authors thank Dr. Jeffey Colgren for valuable comments on the paper, Dr. Carine Le Goff and Alexandre Jan for the phase contrast image of a 1-day old ctenophore. HPF, SBFSEM and TEM imaging was done in the Oxford Brookes Centre for Bioimaging. This work was supported by the Sars Centre core budget.

