## Supplementary Files for "Syncytial nerve net in a ctenophore sheds new light on the early evolution of nervous systems"

**Supplementary Figure 1.** Blebbed neurite morphology of ctenophore SNN neurons

**Supplementary Figure 2.** SNN neuron synapses onto comb cells

**Supplementary Figure 3.** Multiple sensory cells on the epidermis of the ctenophore *M. leidyi*

**Supplementary Table 1.** Sensory cell types detected in *M. leidyi* 1-day old cydippid

**Supplementary Video 1.** 3D reconstruction of the SNN, comb rows, sensory cells, mesogleal neurons and a tentacle from SBFSEM data of a 1-day old cydippid

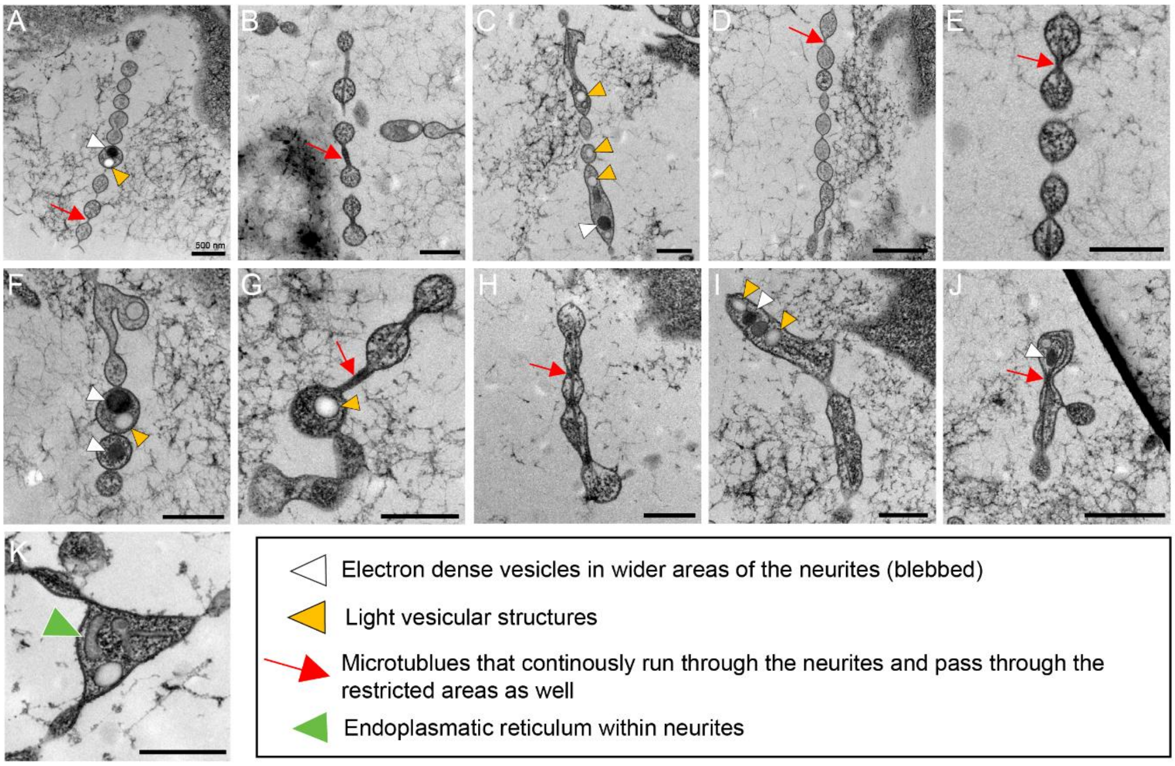


**Supplementary Figure 1. Blebbed neurite morphology of ctenophore SNN neurons.** (A-K) High resolution TEM micrographs of the unique structure of *M. leidyi* SNN neurites. Scale bar in all panels: 500 nm

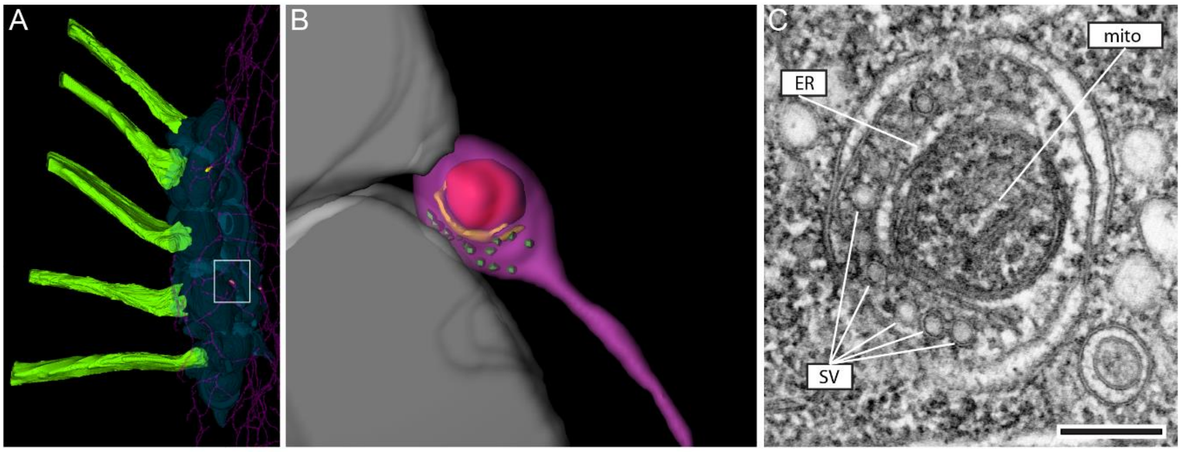


**Supplementary Figure 2. SNN neuron synapses onto comb cells.** (A) From SBFSEM data 3D reconstructed comb row, SNN neurons and 3 synapses (yellow). White box: synapse shown in B. (B) SBFSEM data 3D reconstruction of SNN neuron chemical synapse onto comb cell. Red: mitochondria; yellow: ER; green: synaptic vesicles; grey: comb cells. (C) High resolution TEM micrograph of a chemical synapse contacting comb cells. Note the typical presynaptic triad of mitochondrion, ER and synaptic vesicles. Scale bar: 200 nm

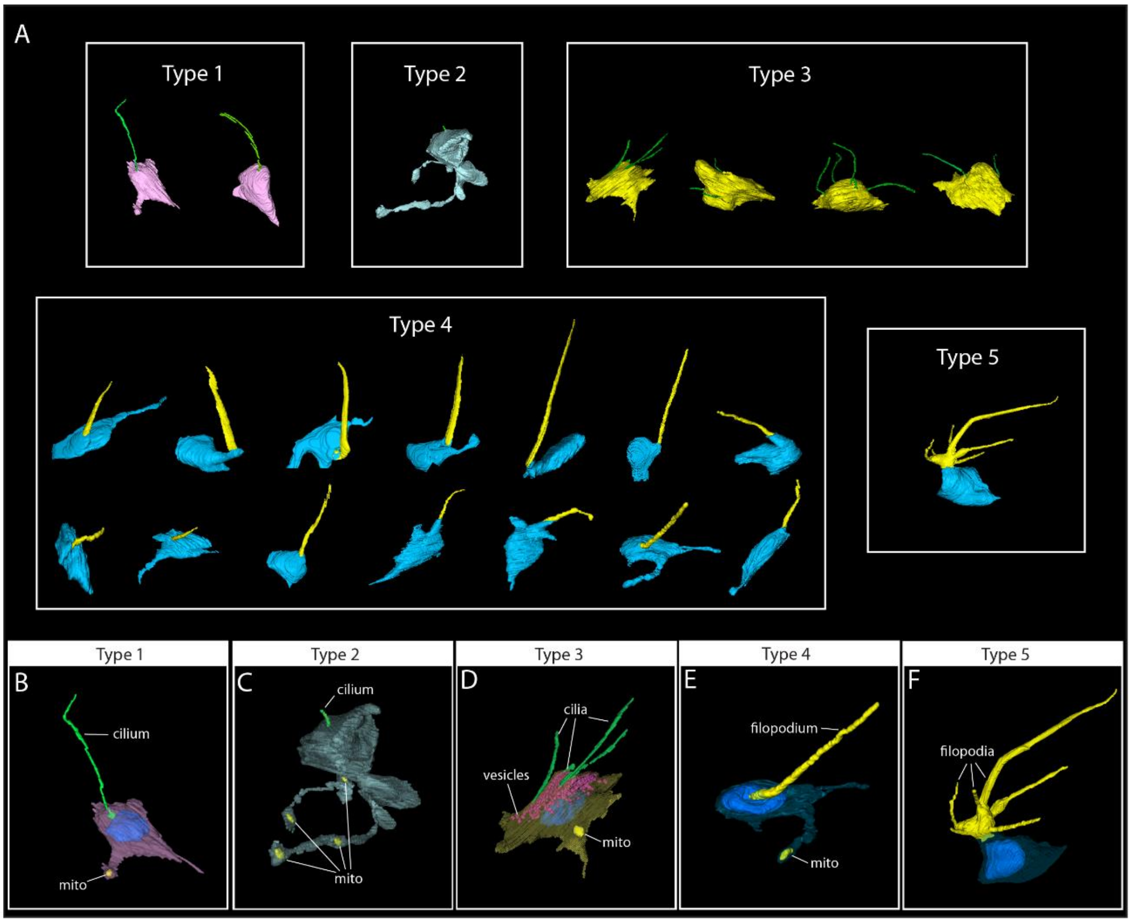


**Supplementary Figure 3. Multiple sensory cells on the epidermis of the ctenophore *M. leidyi*.** (A) 3D reconstructed sensory cells grouped into 5 types based and their cellular protrusions (B-F) Representative sensory cell of each type (1-5).

**Supplementary Table 1. Sensory cell types detected in *M. leidyi* 1-day old cydippid**

| **Sensory cell type** | **Characteristic traits** | **Synapse** |
|---|---|---|
| Type 1 | single long cilium with onion root basal body | yes* |
| Type 2 | single short cilium, no onion root basal body, long neurites | yes* |
| Type 3 | multiple cilia, no onion root basal body, large dense core vesicles underlying cilia | yes* |
| Type 4 | single filopodium | yes* |
| Type 5 | multiple filopodia | no |

*detected in some
